# Selection-free whole genome transplantation revives dead microbes

**DOI:** 10.64898/2026.03.13.711674

**Authors:** Zumra Peksaglam Seidel, Nacyra Assad-Garcia, Vanya Paralanov, Feilun Wu, Olivia Chao, Elizabeth A. Strychalski, Eugenia Romantseva, Tyler Goshia, J. Craig Venter, John I. Glass

**Affiliations:** J. Craig Venter Institute, San Diego, CA 92037, USA; J. Craig Venter Institute, Rockville, MD 20850, USA; National Institute of Standards and Technology, Gaithersburg, MD 20899, USA

## Abstract

We present a living, synthetic bacterial cell made by transplanting a complete genome into a dead cell. After killing *Mycoplasma capricolum* cells by chemically crosslinking their genome with Mitomycin C (MMC), we installed synthetic *Mycoplasma mycoides* genomes into the resulting dead cells using Whole Genome Transplantation (WGT)^1,2^. During WGT, a synthetic donor genome is placed into a recipient cell, thereby reprogramming that cell to adopt a new genetic identity^3^. WGT has only been demonstrated using species within one phylogenetic clade of *Mollicutes* bacteria^4^. A major barrier to expanding WGT to diverse bacterial species has been the inability to inactivate the recipient genome, leading to false positive transplants due to homologous recombination of antibiotic resistance markers from the donor genome into the recipient cell genome. Here, we address this key limitation by removing reliance on an antibiotic resistance marker to select for transplants; recipient cells are dead unless revived by the installation of a new genome. Our work demonstrates a general approach to fully inactivate the recipient cell genome, reports the first living synthetic bacterial cell constructed from non-living parts, and advances WGT for building engineered or synthetic cells for diverse applications.

## MAIN TEXT

While synthetic bacterial cells programmed with synthetic DNA are poised to emerge as enabling biotechnological platforms^5,6^, technical challenges have held back synthetic cells from reaching their full potential. To construct the first synthetic bacterial cell^3^, the JCVI developed three key synthetic biology technologies: (1) we greatly improved the speed and efficiency of assembling synthetic oligonucleotides into DNA molecules >100 kb^7,8^, (2) we used the enormous power of yeast genetics to assemble overlapping DNA segments into complete synthetic bacterial chromosomes as yeast centromeric plasmids (YCPs)^9–11^, and (3) we invented WGT^1,3^ to biomolecularly reprogram the recipient cell by installing a new genome. While *Escherichia coli* strains with synthetic genomes were constructed with some of the methods JCVI developed for synthetic DNA, these strains did not use a yeast intermediary host as for WGT. Instead, they replaced the *E. coli* genome with iterative swapping of (50 to 100) kbp segments in a slow process requiring months^12,13^. This cumbersome approach has not been applied to other synthetic bacteria. In contrast, WGT allows genome replacement in a single step but remains restricted to the *Mycoides* taxon of mycoplasmas, preventing broader application for synthetic cells.

To overcome this limitation, we developed what we term “zombie cells”, inspired by the concept of zombies in the Haitian Vodou tradition, which are bodies revived magically after death^14^. Although no magic was involved, zombie cells are bacterial cells killed by destruction of their native genomes and subsequently revived with a donor genome. In this context, “dead” denotes the loss of capacity for replication. Transplantation of a donor genome into such cells yielded restored viability in a reanimated chassis, effectively rebuilding living cells from non-living components.

In WGT, we install a synthetic version of a *Mycoplasma mycoides* donor genome into a *Mycoplasma capricolum* recipient cell^1,2^ (Figure 1a). Attempts to extend WGT to other bacteria have shown an inverse relationship between the success of transplantation and the phylogenetic distance between donor and recipient cells^4^. If phylogenetic distance were the primary determinant of WGT success, transplantation within a single species should overcome this barrier. However, attempts to extend WGT to intraspecies WGT in *Mycoplasma genitalium*, consistently resulted in false positive transplants despite antibiotic selection (Figure 1b). These outcomes revealed that transplantation attempts involving recipient cell species with active homologous recombination systems—present in most bacteria—yield false positives, as only small genomic regions surrounding antibiotic resistance markers recombine into the recipient genome. Importantly, this complication was not apparent in early WGT studies, because *M. capricolum* lacks an active system for homologous recombination. This allows confident antibiotic-based selection for the donor genome. As WGT is extended to recipient species that retain homologous recombination, antibiotic-based selection becomes unreliable. This highlights the need for a strategy that eliminates reliance on antibiotic markers.

**Figure 1.**
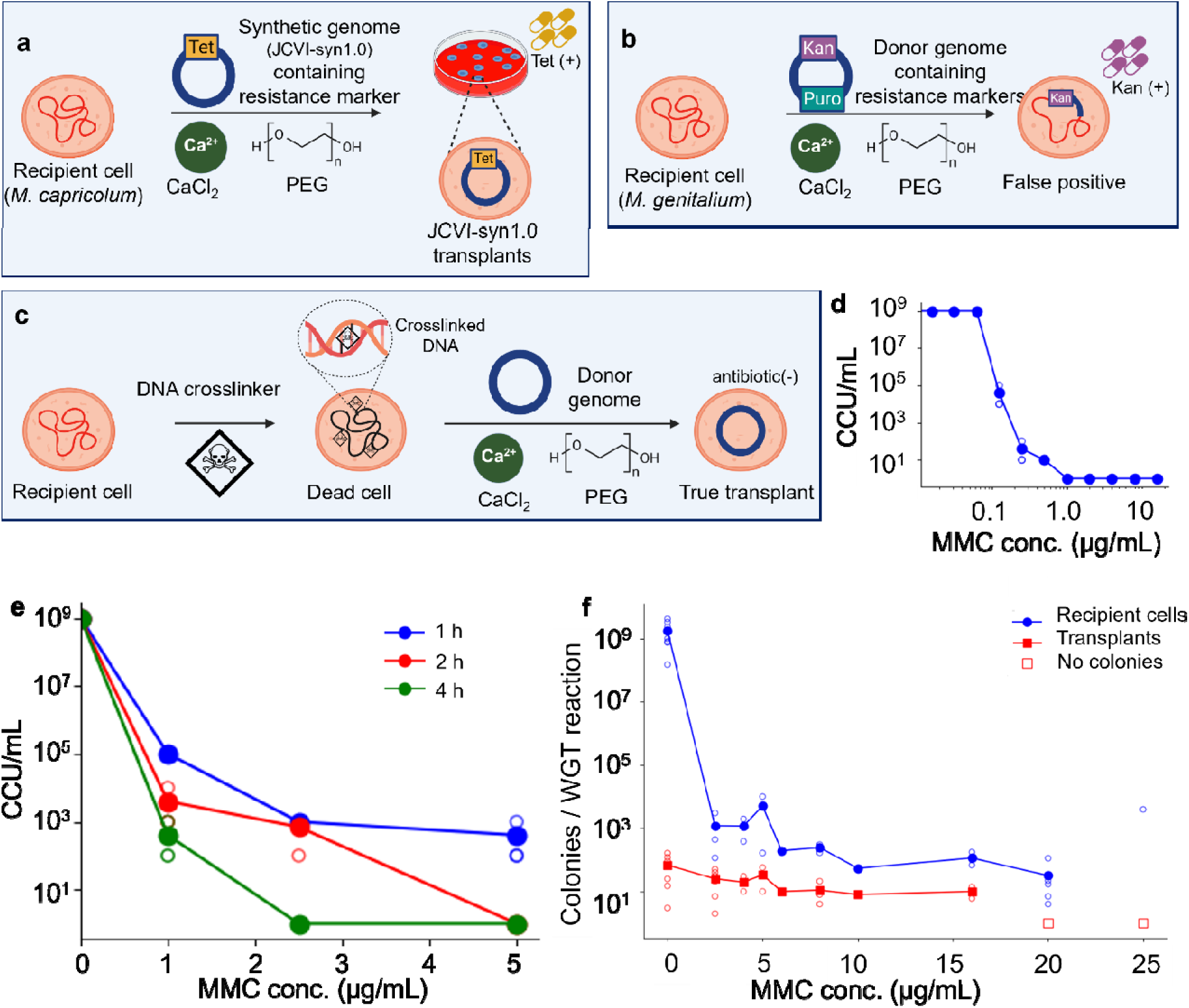
Mitomycin C (MMC) treatment can be incorporated into Whole Genome Transplantation (WGT) to inactivate recipient cells while preserving transplantation capacity. (a) Schematic of the first successful WGT is shown using JCVI-syn1.0 donor genomes containing a tetracycline resistance marker transplanted into *M. capricolum* recipient cells. Following plating on agar plates containing tetracycline, only cells carrying the donor genome survived, yielding JCVI-syn1.0 transplants. (b) Schematic of homologous recombination-driven false-positive transplantation in *M. genitalium* is illustrated. The donor genome contained kanamycin and puromycin resistance markers on opposite sides of the genome. After plating on agar plates containing kanamycin, donor genome regions surrounding kanamycin antibiotic resistance marker recombined into the recipient genome without full genome replacement. These were false positive transplants. (c) Selection free WGT where MMC is added to *M. capricolum* recipient cell cultures in the final hours of growth is diagrammed. The MMC chemically crosslinks the recipient cell genome and kills the cells. Recipient cells that take up a donor genome are revived. (d) The minimum inhibitory concentration (MIC) of *M. capricolum* cells treated with MMC informed our choice of concentrations to investigate for use with WGT. (e) *M. capricolum* cell viability decreased with increasing MMC concentration and treatment duration. (f) The effect of MMC concentration on recipient cell viability and transplantation efficiency is shown. Recipient cell viability decreased by ≈10^6^-fold, whereas transplantation efficiency decreased by ≈10-fold; no transplantation was observed at 20 or 25 µg/mL MMC. In (d) and (e), each data point represents a single CCU plate scored by colour change from red to yellow. In (f), open circles represent a single biological replicate and filled circles indicate arithmetic mean values. Lines are drawn to guide the eye.

Here, we report a general solution to this problem, by killing recipient cells without compromising their capacity for WGT. Specifically, chemically disrupting recipient genomes through DNA crosslinking renders cells genomically inactive, while preserving their capacity to receive and express a transplanted genome. As a proof of principle, we incubated recipient cells in media containing mitomycin C^15^ (MMC), a DNA alkylating agent that irreversibly suppresses genome function. By inactivating viable recipient genomes, we eliminated the need for antibiotic selection of transplants.

This approach has two far-reaching consequences for building synthetic cells. In a conceptual advance, our approach marks the first construction of a living synthetic bacterial cell, revived from the non-living components of a dead cell whose genome we destroyed. In a technological advance, our results overcome an outstanding technical challenge from our 2010 paper^3^ preventing WGT beyond *Mycoides* group *Mollicutes* bacteria. Together, the work presented here promises to finally pair our ability to write synthetic DNA with the ability to place that DNA into a receptive environment for expression.

### MMC treatment of recipient cells

To establish a baseline to compare WGT and selection-free WGT, we first performed WGT as reported previously^1,2^. Donor *M. mycoides* genomes were isolated from living bacteria or *Saccharomyces cerevisiae* cells as yeast centromeric plasmids (YCPs) containing the complete bacterial genome and ∼ 3 kbp yeast vector sequences. Donor genomes were introduced into *M. capricolum* recipient cells, with calcium chloride and polyethylene glycol (PEG), and spread on nutrient agar containing antibiotic to await colony growth. These donor genomes required an antibiotic resistance marker and, often, a *lacZ* gene, allowing selection of transplants by survival and blue-white screening. Only transplants—cells successfully reprogrammed by the donor genome through WGT— survived the antibiotic and yielded blue colonies.

To inactivate the recipient cell genome, we incorporated the DNA crosslinker MMC for selection-free WGT. MMC is an alkylating antibiotic activated by enzymatic reduction of its quinone group, enabling covalent DNA binding and crosslinking at 3′-CpG-5′ sites^15–17^. A single crosslink blocks replication and kills the bacterial cell^15,18,19^. When applied only briefly to recipient cells, MMC crosslinks the genome but leaves transcription and translation machinery intact^16,20^. This allows introduction and expression of the donor genome, while the disabled recipient genome cannot replicate or recombine. While MMC is known to suppress DNA synthesis, higher concentrations can also inhibit RNA and protein synthesis^21^, an effect that we sought to avoid. In this modified workflow (Figure 1c), MMC treatment killed recipient cells, inactivating their genomes before WGT. Transplants were then grown on plates without antibiotics, because only cells containing the donor genome would grow.

To determine the conditions for MMC to kill recipient cells, we chose the *M. capricolum* strain *Mcap*ΔRE as donor genomes isolated from yeast. This strain has its single restriction enzyme gene disrupted, so unmethylated donor genomes isolated from yeast would not be destroyed when transplanted into *M. capricolum*^2^. We measured a minimal inhibitory concentration (MIC) of MMC for *Mcap*ΔRE of 1 µg/mL (Figure 1d) and an inverse relationship between cell viability and both MMC concentration and incubation time (Figure 1e). Based on these results, we chose a 2 h incubation and MMC concentrations between (0 to 25) µg/mL for selection-free WGT.

We quantified both the number of recipient cells that survived exposure to MMC and transplants by counting the resulting colonies after MMC treatment (Figure 1f). Surviving *Mcap*ΔRE recipient cells produced white colonies, while a l*acZ* gene in the donor genome of transplants produced blue colonies. Additionally, recipient cell viability dropped by ≈ 10^6^, whereas transplantation decreased only ≈ 10-fold. Although transplantation occurred at MMC concentrations up to 16 µg/mL, we observed no transplant colonies at higher concentrations, suggesting the onset of inhibition of RNA synthesis, protein translation, or both. We therefore further tuned MMC treatment of recipient cells prior to WGT to 2.5 µg/mL MMC for 2 h to effectively disable the recipient genome, while preserving cellular functions necessary for donor genome expression. Thus, we achieved selection-free WGT by applying MMC at low doses to crosslink DNA.

### Transplantation into Dead Cells

Treatment with MMC significantly improved the number of transplants relative to the number of viable recipient cells. (Figure 1f). Taking this quantity as a metric for efficiency, we found a ≈ 500,000-fold increase in efficiency for selection-free WGT protocol. Specifically, WGT without MMC began with ≈ 10^10^ recipient cells and yielded 68 ± 59 transplants (N = 7; Methods), corresponding to one transplant per ≈ 1.5 × 10^8^ viable recipient cells. Selection-free WGT with MMC caused viable recipient cells to drop dramatically to ≈7.2 × 10^3^ cells, while still yielding 25 ± 19 transplants per reaction (N = 6; Methods), corresponding to one transplant per ≈ 288 recipient cells. These results imply either MMC treatment rendered the remaining viable cells extraordinarily competent for WGT, or we achieved our goal of transplantation into dead recipient cells. Consideration of the original development and assumptions underlying WGT favor the latter interpretation.

Previous demonstrations of WGT used *M. capricolum* recipient cells optimized using transformation assays with a replicative plasmid, assuming the parameters affecting transformation also influence transplantation. We hypothesized that if MMC treatment rendered recipient cells more amenable to transplantation, those same cells should also show increased transformation. With identical MMC treatments, transplantation continued to occur (Figure 2c, 2d), while no transformation was observed. Because transformation requires functional replication (Figure 2a, 2b), this provided evidence for viable recipient cells. These results therefore supported the conclusion that donor genomes were inserted into dead recipient cells and expressed to revive and genomically reprogram the cells.

**Figure 2.**
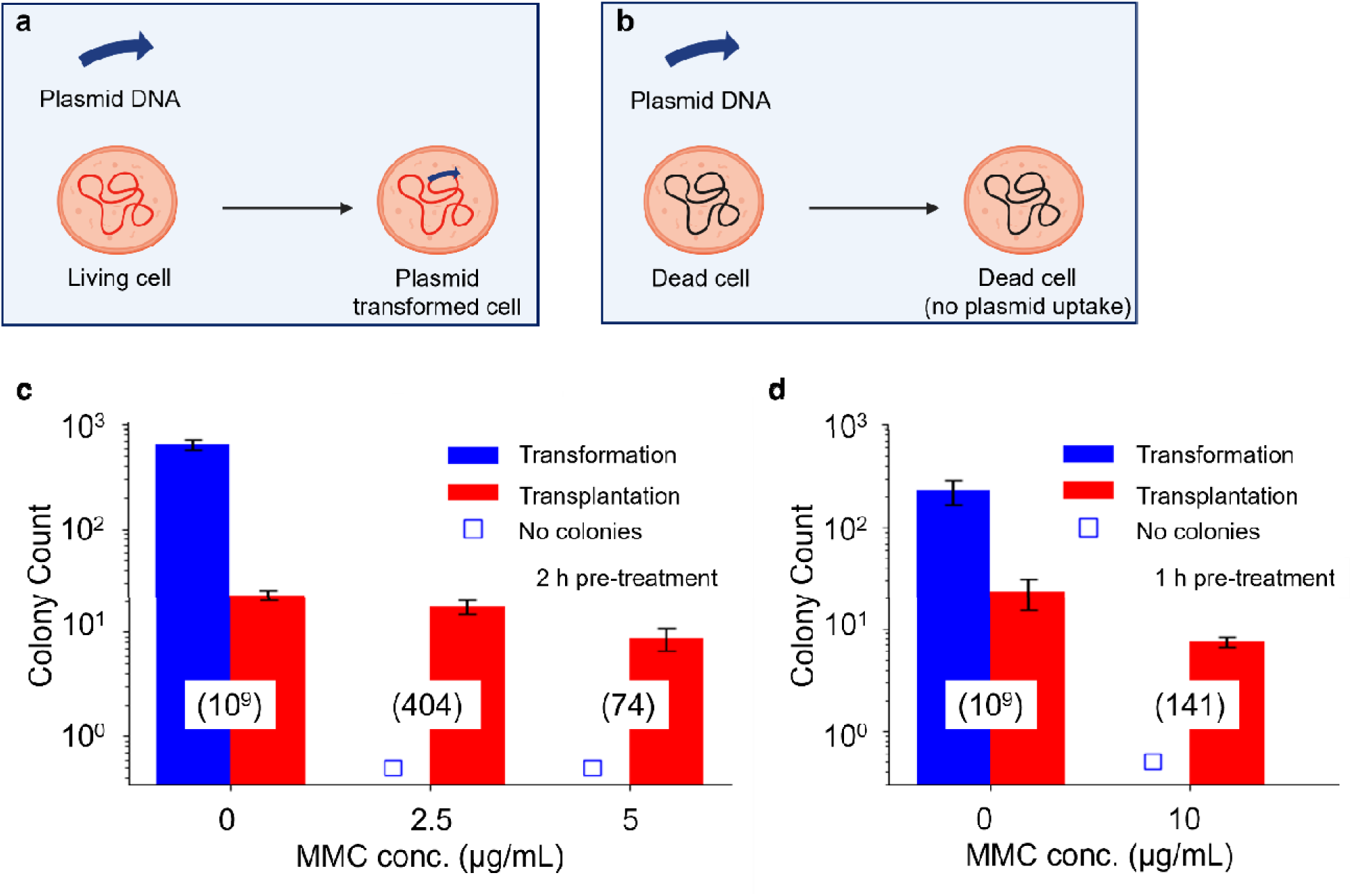
MMC treatment enables WGT into dead recipient cells. The schematics compare replicative plasmid transformation in (a) living recipient cells (b) dead recipient cells. Genome-inactivated cells did not support plasmid transformation. Plasmid transformation and genome transplantation efficiencies were compared following MMC treatment. Results from recipient cells treated with 2.5 µg/mL or 5 µg/mL MMC for 2 h (c) and 10 µg/mL MMC for 1 h (d) provide evidence for transplantation into dead cells. Numbers in parentheses indicate viable recipient cell counts for each condition.

### Selection-free WGT

Previous WGT protocols relied on tetracycline selection to allow the growth of only transplants expressing JCVI-syn1.0 donor genomes, which carry a *tetM* marker (Figure 3a, top left). This selection was necessary, because recipient cells without donor genomes grew to form a lawn on plates without antibiotic, turning the plate from red to yellow due to a pH shift detected by phenol red (Figure 3a, bottom left, inset). Plating dilute cultures of recipient cells resulted in distinct white colonies, while blue colonies from transplants no longer appeared. However, selection-free WGT grew blue transplant colonies on plates without antibiotic (Figure 3a, top and bottom right). This indicated that inactivation of the recipient genome by a DNA crosslinking agent eliminated the need for selection, as recipient cell genome was suppressed.

**Figure 3.**
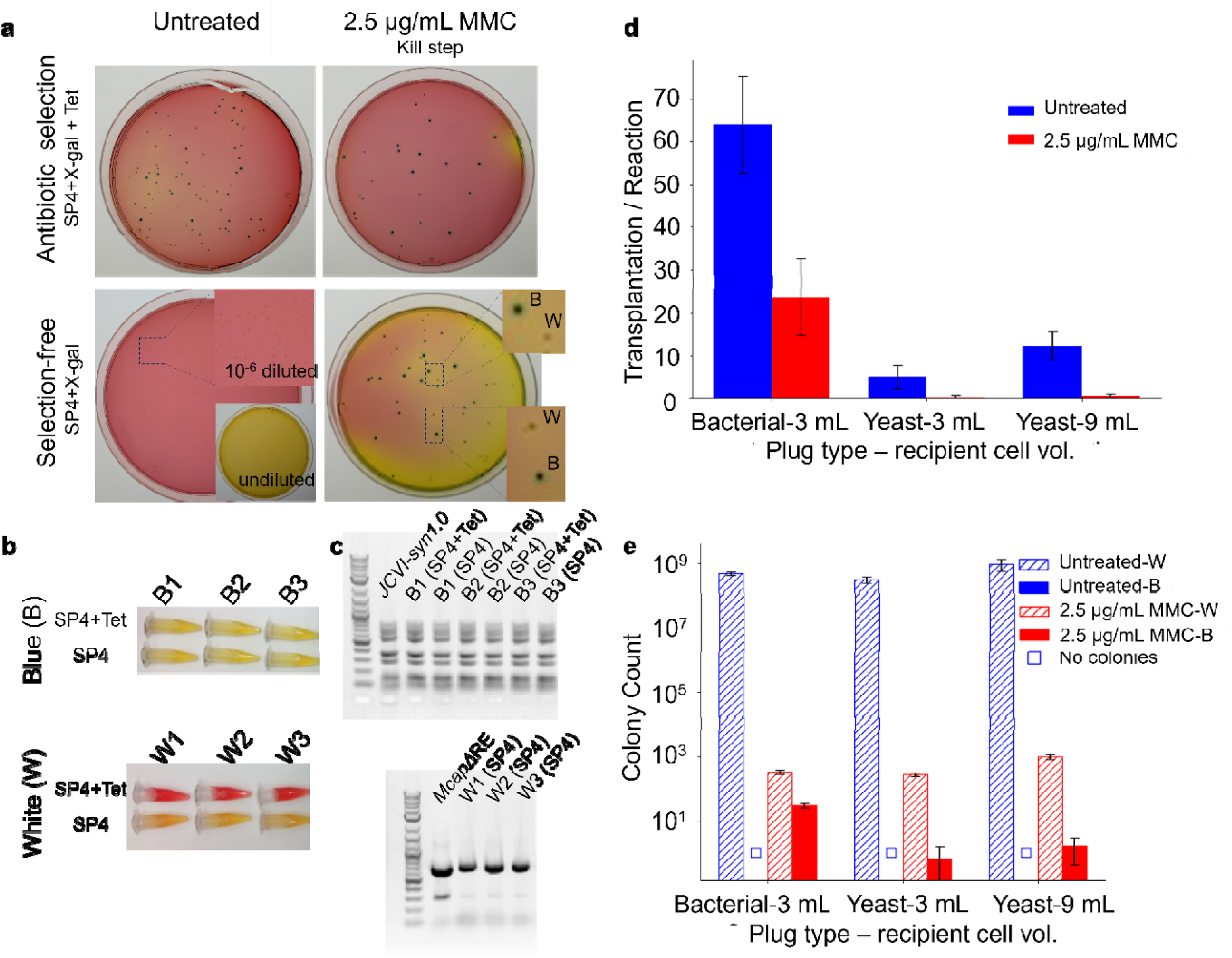
MMC treatment during WGT obviates the need for antibiotic selection of transplants. We compared the efficiency of WGT with and without antibiotic selection in the presence and absence of MMC treatment. (a) Representative photographs of 10-cm nutrient agar plates show white (*M. capricolum*) and blue (JCVI-syn1.0) colonies after (8 to 16) h growth in SP4 medium with and without tetracycline. The JCVI-syn1.0 genome contains a tetracycline resistance marker, and it can grow in both in SP4 medium with and without tetracycline, whereas *M. capricolum* grows only grows in SP4 without antibiotic. (b) We inoculated 1.8 ml tubes containing SP4 growth media with and without tetracycline with cells taken from three blue and three white colonies. After 3 days of incubation at 37 °C, cell growth was indicated by increased acidity causing the phenol red in the media to change from red to yellow. (c) Additional confirmation of colony genotype was done using multiplex PCR of the aforementioned sets of white and blue colonies. Experiments (b) and (c) confirmed white colonies were *M. capricolum* and blue colonies were JCVI-syn1.0 transplants. (d) Transplantation efficiency was measured for untreated and MMC-treated recipient cells using bacterial or yeast genome plugs across varying recipient cell volumes. (e) The numbers of white and blue colonies on antibiotic-free plates from (d) were compared with untreated cells plated at a 10^-6^-fold dilution to prevent lawn formation, as shown in the bottom panel of (a). Because this dilution is too low to see individual colonies, successful transplants were not observed in untreated cells, making selection-free plating impractical for WGT without inactivating genome of recipient cells. In (d) and (e), bar graphs show arithmetic mean values, and black error bars represent standard deviation from three biological replicates.

Although most recipient cells were killed by MMC treatment, a few remained viable. When grown on plates without antibiotic, both blue and white colonies were observed (Figure 3a bottom right). To confirm viable transplants and living recipient cells, individual blue and white colonies were picked and grown separately in SP4 medium with and without tetracycline (Figure 3b). White colonies grew only in SP4 but not in SP4+Tet, confirming these as recipient *Mcap*ΔRE cells that survived MMC treatment and did not undergo transplantation. Blue colonies grew in both conditions, verifying these cells as successful transplants carrying the *tetM* marker of JCVI-syn1.0 donor genomes. Growth was indicated by the colour shift of the growth medium from red to yellow, reflecting acidification associated with cell metabolism and consistent with our observations of growth on agar plates.

We performed multiplex polymerase chain reaction (PCR) analyses to further confirm WGT and selection-free WGT yield blue colonies with JCVI-syn1.0 genomes and similar numbers of transplants per reaction. Individual blue and white colonies shown in Figure 3b served as templates. Nine amplicon pairs, along with yeast rRNA and His3 primer pairs (Table 1), allowed identification of the JCVI-syn1.0 synthetic genome. All of these 11 expected bands were detected in blue colonies grown in either SP4 or SP4+Tet, with band intensities matching the JCVI-syn1.0 control, confirming blue colonies as transplants carrying the complete JCVI-syn1.0 genome (Figure 3c). For white colonies grown on SP4 agar, we observed one bright band and three faint bands, characteristic of the *Mcap*ΔRE control. This provides evidence white colonies grew from recipient cells that survived MMC treatment without undergoing transplantation.

**Table 1.**
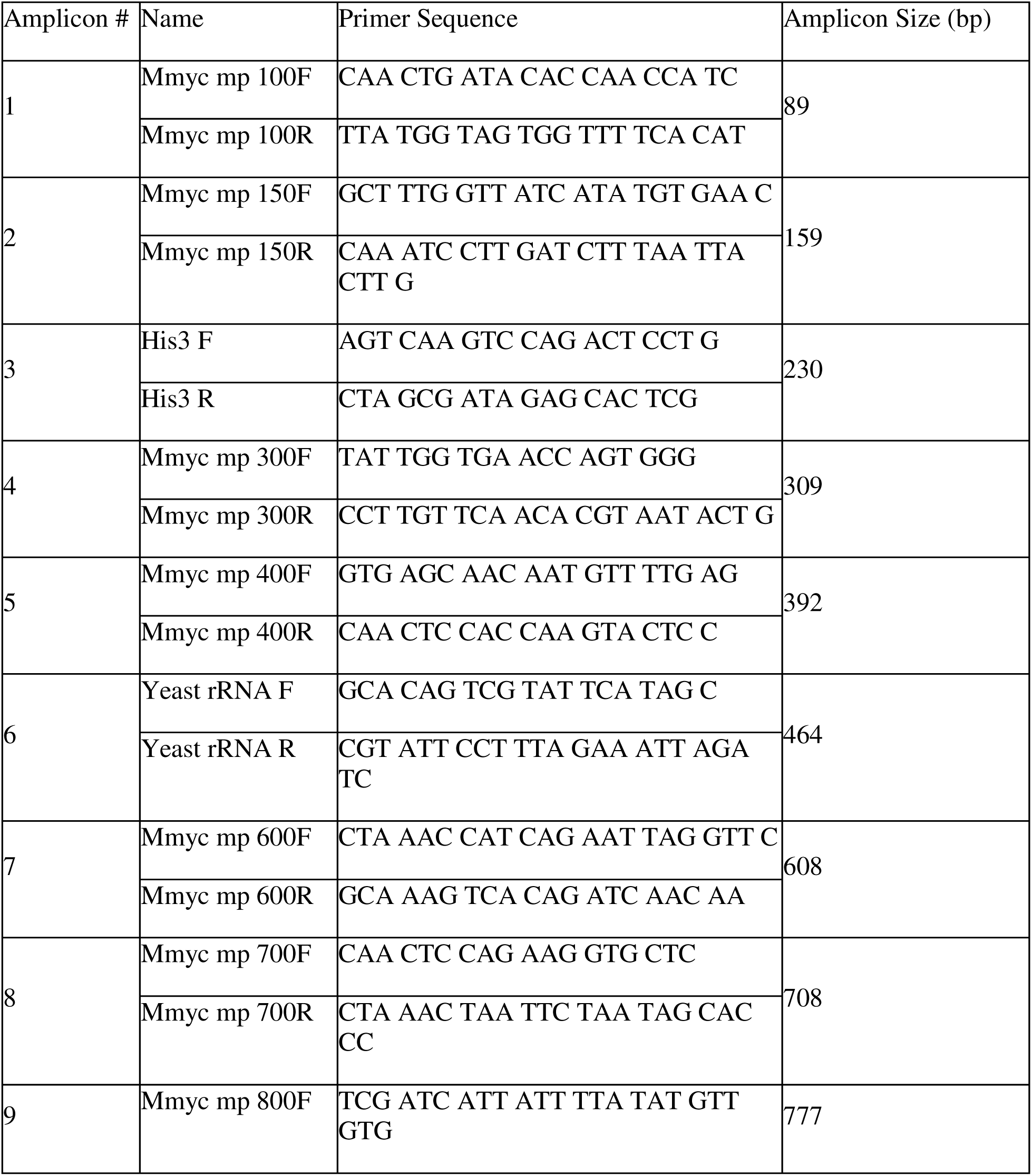
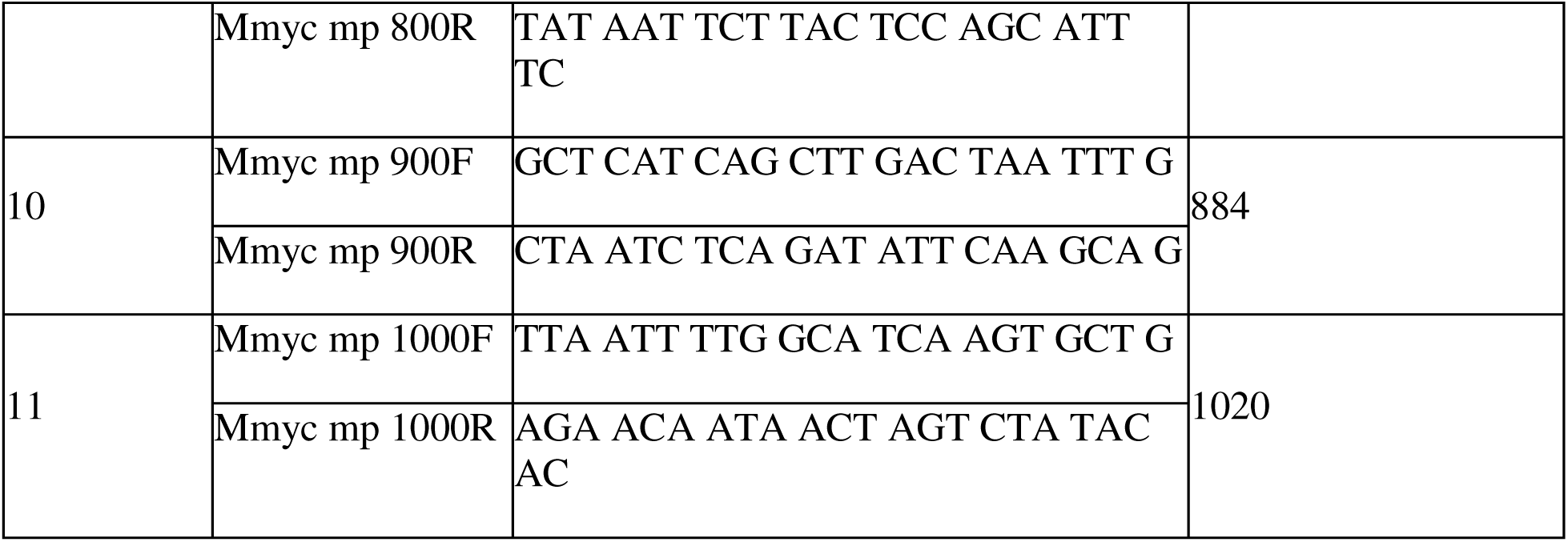
Multiplex PCR amplicons used to identify JCVI-syn1.0 genome.

We next investigated selection-free WGT using donor genomes prepared from bacterial or yeast sources using various input numbers of recipient cells (Figure 3d-e; Methods). Bacterial genome plugs yielded 64 ± 14 transplants (N = 3; Methods) using WGT, whereas selection-free WGT yielded 24 ± 11 transplants (N = 3; Methods and Figure 1d). In contrast, WGT from yeast plugs was consistently less efficient, yielding 5 ± 3 and 12 ± 4 transplants (N = 3) from original WGT for 3 mL and 9 mL recipient cell volumes, respectively, and only 1 transplant (N = 3) for selection-free WGT. As reported previously^3,4^, transplantation efficiency was higher using bacterial genome plugs than yeast genome plugs for both WGT protocols. These results demonstrate transplantation using selection-free WGT and donor genomes prepared using either bacteria or yeast and still allowed WGT into MMC killed *Mcap*ΔRE cells.

### Crosslinking DNA Generates Zombie Cells

To test whether DNA crosslinking of recipient genomes provides a general strategy for generating zombie cells, we examined an alternate chemical to induce replication-blocking DNA lesions prior to WGT. MMC is activated *in vivo* through reductive enzymes that convert its quinone into a highly reactive alkylating agent, with subsequent loss of its methoxy group. Once activated, MMC forms monoalkylated adducts and inter-strand or intra-strand DNA crosslinks at guanine, with a strong preference for 5’-CpG-3’ sequences^15^. These crosslinks block replication and repair processes, driving genome inactivation (Figure S3). Psoralen, another DNA crosslinking compound, intercalates between DNA base pairs due to its small, planar structure. Upon photoactivation with ultraviolet A (UVA) radiation (315–400 nm)**, it** forms mono- and bi-functional adducts with thymine bases at 5’-TpA-3’ sequences^22^ (Figure S4), effectively blocking replication and enabling light-controlled genome inactivation, whereas psoralen or UVA alone have minimal effect on cell viability^23^.

Figure 4a compares psoralen with 30 min UVA treatment with MMC treatment, to evaluate alternative crosslinkers. In contrast to MMC, psoralen was significantly more cytotoxic and supported WGT only within a narrow range of concentration, yielding ≈ 10 % WGT efficiency. Increasing UVA exposure from 30 min to 60 min further reduced the number of transplants (Figure 4b). Thus, selection-free WGT is not limited to MMC and requires the ability to selectively disable recipient genome replication while maintaining an otherwise functional recipient cell. Together, these findings demonstrate that recipient genome inactivation with DNA crosslinker constitutes a modular extension of WGT that supports selection-free transplantation, mitigates homologous recombination associated false positives and enables the use of donor genomes without antibiotic resistance markers.

**Figure 4.**
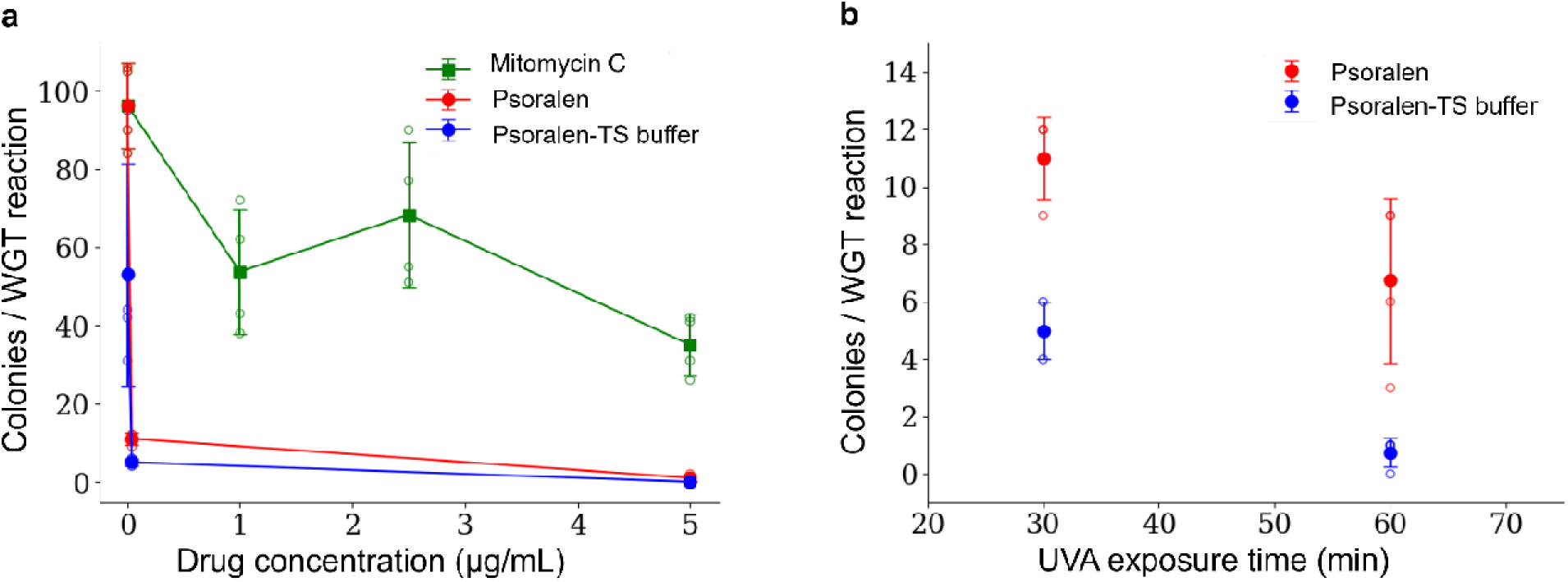
Alternative DNA crosslinking agents support selection-free WGT. (a) Transplantation efficiency was compared in MMC and psoralen treated recipient cells. (b) Increasing UVA exposure time following treatment with light-activated psoralen at 0.04 µg/mL lowers transplantation efficiency. In (a) and (b), open circles represent a single biological replicate and filled circles indicate arithmetic mean values. Lines are drawn to guide the eye.

## Discussion

Since the first demonstration of WGT, we have been working to clarify its molecular mechanisms^2,4,10^ and enable its application beyond a small set of mycoplasmas^24,25^. Here, selection-free WGT using dead recipient cells with a chemical crosslinked genome circumvented the challenge posed by homologous recombination that prevented intra-species WGT in non-*Mycoides* group *Mollicutes* bacteria.

We investigated the DNA crosslinking agents MMC and psoralen preparing zombie cells, which lack replication but preserve membrane and metabolic function. Several other strategies also be used to inactivate the recipient genome; however, they can fail to maintain a viable, transcriptionally and transitionally active recipient chassis. For example, nitrogen mustards non-selectively alkylate DNA, proteins, and membrane lipids. Similarly, platinum-based drugs cause severe helix distortion and global cellular stress^26^. On the other hand, clustered regularly interspaced short palindromic repeats-mediated interference (CRISPRi) systems^27^ potentially offer another approach. CRISPRi could be introduced into WGT for inducible and targeted silencing of genes associated with homologous recombination prior to transplantation. This would circumvent antibiotic selection. The CRISPRi strategy may also offer a superior approach to zombie cells, because CRISPRi would not risk inactivation of ribosomal RNA by MMC. There could be other strategies to advance WGT that overcome homologous recombination problem.

Despite our progress here, the extension of WGT beyond the *Mycoides* taxon of mycoplasmas faces additional hurdles. For instance, a little-known attribute of our *M. capricolum* recipient cells is the absence of a calcium-activated DNA endonucleases on the cell surfaces. Almost all bacteria have endonuclease activity.^28^. In other bacteria, extracellular DNA is degraded rapidly in the presence of calcium chloride. This cleavage of the donor DNA would prevent WGT with existing protocols. We actively seek strategies to overcome this limitation, and additional species-specific barriers may yet emerge. Ultimately, developing fit-for-purpose recipient strains, disruption of the recipient genome, and intraspecies transplantation strategies may all provide leverage to expand the utility of WGT and enable synthetic cells.

### Conclusion & Future Work

Importantly, we report here construction of a living bacterial cell using non-living parts. This stands as a milestone for building synthetic cells. We accomplished this by altering WGT to install the donor genome into a genome-inactivated “dead” recipient cell rather than a living recipient cell. This overcomes challenges related to homologous recombination and achieves an outstanding task needed for generalizing WGT beyond the *Mycoides* taxon of mycoplasmas. By inactivating the recipient genome while preserving cellular function, we show the revival of a non-living cell by a transplanted synthetic genome. Here, we demonstrated our zombie cell-WGT approach to construct living bacterial cell from non-living parts. This strategy highlights that the minimal requirement for “booting up” a cell may only require intact transcription and translation machinery, rather than a fully living recipient, opening new avenues for exploring the fundamental principles of cellular life.

The use of such zombie cells redefines dead *versus* living as a distinction between genome replication rather than cellular viability. Our findings suggest that naturally-evolved biology routinely may operate across a porous boundary between life and death, and this boundary can now be intentional and engineerable. As remaining challenges are addressed, WGT will offer a general platform for installing synthetic genomes into engineered cells, accelerating the development of synthetic cells for biotechnology, medicine, environmental applications, and beyond.

## Methods

### Microbial Strains

*Mycoplasma capricolum* lacking restriction enzymes (*Mcap*ΔRE) was used as the recipient cell^2^. Donor genomes were obtained from *Mycoplasma mycoides* JCVI-syn1.0 bacteria cells or as a yeast centromeric plasmid^3^. For transformation experiments, pMYCO1 plasmid^29^ was used.

### Growth Media

*Mcap*ΔRE recipient cells were grown in SSG medium prior to WGT^10^. Recipient cells were recovered post translation in SP4 medium supplemented with 17% (v/v) heat-inactivated foetal bovine serum (Gibco, Cat. No. 16140-071), referred to as SP4+FBS. JCVI-syn1.0 donor cells were inoculated and cultivated in SP4 medium supplemented with 17% (v/v) KnockOut™ serum replacement (Fisher Scientific, 10828028), referred to as SP4+KO. During WGT, donor genomes and recipient cells were incubated in SP4 media without serum, where an equal volume of distilled deionized water was substituted for the serum component^1,2^.

Growth plates to screen for *LacZ*-expressing transplants based on colony colour contained SP4, 17% horse serum (Gibco, 26050-088), 1 % Noble agar (BD, 214230), and 150 µg/mL X-gal (Goldbio, 7240-90-6). Growth plates for antibiotic selection in *Mcap*ΔRE recipient cells using WGT contained SP4, 17% horse serum (Gibco, 26050-088), 1 % Noble agar (BD, 214230), 4 µg/mL tetracycline (Sigma-Aldrich, T7660), and 150 µg/mL X-gal. Growth plates for selection-free WGT contained SP4, 17% horse serum (Gibco, 26050-088), 1 % Noble agar (BD, 214230), and 150 µg/mL X-gal. Growth plates for measurements of colony forming units (CFU) to enumerate *Mcap*ΔRE recipient cells contained SP4, 17% horse serum (Gibco, 26050-088), and 1 % Noble agar (BD, 214230).

### Colour Changing Unit (CCU) assay

Twelve 10-fold serial dilutions of *Mcap*ΔRE cells were made in SP4+KO medium and incubated at 37 °C for three days. Acid produced by growing cells turn the phenol red indicator in the media from red to yellow. Bacterial concentration as CCU/mL was calculated from the most dilute sample that exhibited a colour change visible by eye ^30^.

### MMC Minimum Inhibitory Concentration (MIC)

To perform the MIC assay, *Mcap*ΔRE cells were grown in 5 mL of SP4+KO at 37 °C to pH 6.3 ± 0.1 or ≈ 10^9^ CCU/mL. A 16 µg/mL MMC working solution was prepared by adding MMC stock solution (Sigma, M5353; 10 mg/mL in DMSO) to 15 mL SP4+KO. MMC treatment was performed in 96-well plates as follows: 360 µL of 16 µg/mL MMC working solution added to wells A12 to H12 and 180 µL of SP4+KO to the remaining wells. A two-fold serial dilution of MMC was performed by transferring 180 µL from A12–H12 to A11–H11, mixing 3–5 times, and continuing the dilution stepwise to A2–H2, discarding 180 µL from the final wells. Wells A1–H1 served as antibiotic-free controls. 20 µL of *Mcap*ΔRE cells were added (per well, A1–A12), followed by 10-fold serial dilutions down each column (A→H) using a multichannel pipette. To prevent cross-contamination, new pipette tips were used for each dilution step. Three replicate MMC plates were prepared, sealed with parafilm (Parafilm M, PM999), and incubated at 37 °C for (48 to 72) h until no further colour change was observed by eye. The MIC was recorded as the lowest MMC concentration at which no colour change occurred in the corresponding 10-fold dilution, i.e., the first well showing no visible growth.

### MMC Concentration and Incubation Time

To determine suitable conditions for MMC treatment for selection-free WGT, four concentrations of MMC were tested, 0 µg/mL, 1 µg/mL, 2.5 µg/mL, and 5 µg/mL, along with 3 incubation times, 1 h, 2 h, and 4 h. A 100 µg/mL MMC working solution was prepared by diluting MMC stock solution into SP4+KO medium; this working solution was used to prepare 1 µg/mL, 2.5 µg/mL, and 5 µg/mL MMC in SP4+KO medium. A control MMC solution was prepared by mixing DMSO in SP4+KO medium; corresponding to 0 µg/mL MMC.

*Mcap*ΔRE cells were grown in 20 mL of SP4+KO medium at 30 °C to pH 6.3±0.1 or ≈ 10^9^ CCU/mL. The culture was divided into twelve 1 mL aliquots and incubated for 1 h, 2 h, and 4 h at 37 °C. For each MMC treatment, 50 µL of the MMC (0 µg/mL, 1 µg/mL, 2.5 µg/mL, and 5 µg/mL) was added to 1 mL of cells, ensuring equal DMSO concentration in all samples. During incubation, 96-well plates were prepared with 180 µL of SP4+KO medium per well. At each time point (1 h, 2 h, and 4 h), 20 µL of treated cells were added to triplicate wells as follows: A1 to A3 (control), A4 to A6 (1 µg/mL MMC), A7 to A9 (2.5 µg/mL MMC), and A10 to A12 (5 µg/mL MMC). Ten-fold serial dilutions were performed from rows A to H using a multichannel pipette, mixing 4 to 5 times per step, and changing tips between rows to prevent cross-contamination. Plates were sealed with parafilm and incubated at 37 °C for (48 to 72) h until no further colour change was observed by eye.

### Preparation of Agarose Plugs containing mycoplasma

Bacterial agarose plugs containing the JCVI-syn1.0 genome were prepared as described previously^1^, substituting chloramphenicol for tetracycline and streptomycin, and performing lysis and washing steps individually for each plug. Briefly, JCVI-syn1.0 mycoplasma cultures (50 mL) were grown in SP4+KO medium at 37 °C to pH 6.4 ± 0.1. 50 µg/mL chloramphenicol was added, followed by incubation at 37 °C for 90 min. Cells were pelleted by centrifugation (4,000 × g, 15 min, 10 °C), washed in 10 mL Tris–Sucrose buffer (pH 6.5), and centrifuged again under the same conditions. The final pellet was resuspended in 500 µL of Tris–Sucrose buffer. In parallel, 1 mL of 2% low-melting-point (LMP) agarose was melted at 65 °C and maintained at 50 °C. The resuspended cell pellet was equilibrated at 50 °C and mixed at a 1:1 ratio with molten agarose by gentle pipetting. Approximately 90 µL of the agarose–cell mixture was dispensed into each plug mold (BioRad, 170-3591; CHEF Mammalian Genomic DNA Plug Kit) and allowed to solidify at room temperature, yielding ≈10 plugs. After solidifying, each plug was transferred to an individual 1.7 mL microcentrifuge tube and incubated with 312 µL of the Proteinase K solution (prepared from 3000 µL Proteinase K reaction buffer and 120 µL of Proteinase K stock (20 mg/mL Proteinase K, 10 µM CaCl_2_, 50 mM Tris-HCl, 25% glycerol, pH 6.5)) at 50 °C for at least 24 h. The plugs were then washed four times with 1 mL of Tris–EDTA buffer A (20 mM Tris-HCl, 50 mM EDTA, pH 8.0) for 1 h each on a tube rocker. After the final wash, 1 mL of Tris–EDTA buffer A was added, and the plugs were stored at 4 °C until use.

### Preparation of Agarose Plugs Containing Yeast

Yeast agarose plugs containing the JCVI-syn1.0 genome were prepared following previously described protocols^3,10^, performing lysis and washing steps individually for each plug. Briefly, *Saccharomyces cerevisiae* cultures (50 mL) were grown in Complete Minimal medium (Teknova, C8112) at 30 °C with shaking at 220 rpm to an OD_600_ of 1.5–2.0. Cells were pelleted by centrifugation (4,000 × g, 5 min, room temperature), washed once with 25 mL sterile ultrapure water (18.2 MΩ-cm resistivity), and again with 50 mM EDTA (pH 8.0). The pellet was resuspended in 150 µL of cell resuspension buffer (BioRad, CHEF Mammalian Genomic DNA Plug Kit), and mixed with 85 µL of Zymolyase-100T solution (20 mg/mL Zymolyase-100T, US Biological, Cat. No. Z1004, 50% glycerol, 20% glucose, 50 µmol/L Tris) and an equal volume of 2% LMP agarose. Approximately 90 µL of the mixture was dispensed into each well of plug molds and allowed to solidify at room temperature, yielding 6 plugs. After solidifying, each plug was transferred to an individual 1.8 mL microcentrifuge tube and incubated with 183 µL of lytic enzyme solution (prepared from 600 µL Lyticase buffer and 500 µL Zymolyase-100T stock) at 37 °C for 2 h. After this step, the yeast agarose plugs were processed the same as bacterial agarose plugs.

### Release of donor genomes

Agarose plugs were washed three times with 1 mL of Tris–EDTA buffer B (10 mM Tris-HCl, 1 mM EDTA, pH 8.0) for 30 min each on a tube rocker at room temperature. After the final wash, the buffer was completely removed by pipetting. Each plug was digested in 10 µL of 10× β-Agarase buffer, incubated at 42 °C for 10 min, at 65 °C for 10 min, then cooled to 42 °C for 10 min. Digested plugs were incubated at 42 °C for 1 h with 3 units of β-Agarase I (NEB, M0392) per plug.

### WGT

Transplantation was performed as described previously^10^ with 2 h post-transplantation recovery at 37 °C^1,3^. These experiments were done by different authors at different times and JCVI laboratories. Because transplantation efficiency depends strongly on the operator and instrumentation, parameters such as recipient cell volume, recovery duration, and centrifuge speed were optimized individually for each experiment. For clarity, all transplantation procedures are described in terms of a single recipient cell volume for each experiment. In practice, recipient cells were processed in bulk during the initial centrifugation, washing, and CaCl_2_ incubation steps. Following incubation with CaCl_2_, the cell suspension was divided, and individual WGT experiments were initiated in separate 10 mL tubes.

For WGT, *Mcap*ΔRE recipient cells were grown at 30 °C in SSG medium to pH 6.3 ± 0.1. Either 6 mL or 12 mL of cell culture was used for WGT with mycoplasma or yeast plugs, respectively. Cells were pelleted by centrifugation (4000 × *g*, 15 min, 10 °C), washed with 3 mL (mycoplasma) or 6 mL (yeast) of wash buffer (10 mmol/L Tris-HCl, 25 mmol/L NaCl, pH 6.5), and centrifuged again (4000 × *g*, 15 min, 10 °C). The cell pellet was resuspended in 200 µL (mycoplasma) or 400 µL (yeast) of 0.1 M CaCl_2_ and kept on ice for 30 min. In parallel the donor genome mixture was prepared in a 10 mL tube by combining 400 µL (mycoplasma) or 800 µL (yeast) of SP4 without serum with 20 µL (mycoplasma) or 100 µL (yeast) of digested plug. After incubation in CaCl_2_, the recipient cell suspension was added to the donor genome mixture by gentle pipetting, followed by 620 µL (mycoplasma) or 1300 µL (yeast) of 2× fusion buffer (20 mM Tris-HCl, 250 mM NaCl, 20 mM MgCl_2_, 10% PEG_6000_, pH 6.5). These WGT reactions were incubated at 30 °C for 90 min, then supplemented with 5 mL SP4+FBS (30 °C) and allowed to recover at 37 °C for 2 h. Reactions were then centrifuged (4000 × *g*, 15 min, room temperature), resuspended in SP4+FBS, and plated on SP4 agar containing tetracycline and X-gal. Plates were incubated at 37 °C for (2 to3) days, and blue transplant colonies were counted for experimental analysis. Transplant numbers are reported as mean value ± standard deviation from independent biological replicates (N indicated in figure legends) throughout this paper.

### Selection-free WGT

*Mcap*ΔRE cells were grown in SSG medium to pH 6.7 ± 0.1, then treated with (2.5 to 25) µg/mL of MMC, and grown for an additional 2 h at 30 °C to pH 6.3 ± 0.1. The protocol then proceeded the same as for WGT.

### Effect of MMC Concentration on WGT

To assess the effect of MMC treatment on WGT efficiency, *Mcap*ΔRE recipient cells were grown in SSG medium at 30 °C to pH 6.7 ± 0.1. For each WGT reaction, MMC (10 mg/mL) was added to 3 mL *Mcap*ΔRE recipient cells to a final concentration of (0 to 25) µg/mL. Cells were incubated at 30 °C for 2 h, after which 10 µL aliquots were taken for CFU determination. WGT was then performed as described above with centrifugation at 4575 ×*g*, using 20 µL of digested mycoplasma plugs, and reactions were plated directly on SP4 agar plates with tetracycline without a recovery time.

### Effect of MMC on WGT and Transformation

To compare the effects on MMC on WGT and transformation, *Mcap*ΔRE recipient cells were grown to pH 6.7 ± 0.1. For each reaction, 3 mL of cells were collected, and (2.5, 5, and 10) µg/mL MMC was added. Cultures treated with (2.5 and 5) µg/mL MMC and their respective controls were incubated 2 h at 30 °C; cultures treated with 10 µg/mL MMC and its controls were incubated 1 h at 30 °C. WGT was performed as described above using centrifugation at 4575 ×*g* and 20 µL of digested mycoplasma plugs. In parallel, transformation of the pMYCO1 plasmid^29^ was carried out under identical conditions, substituting 1 µg of pMYCO1 plasmid for mycoplasma DNA plugs. Following final resuspension, cells were plated without a recovery period on SP4 agar plates with tetracycline.

### Selection-free WGT with Psoralen, a UV-Activated DNA Crosslinker

To evaluate the effects of different DNA crosslinking agents on WGT efficiency, MMC treatment was compared with treatment with psoralen (4′-aminomethyltrioxsalen; Thermo J60042-MC), a UV-responsive DNA crosslinking compound. The Tris-HCl wash buffer (10 mmol/L Tris-HCl, 25 mmol/L NaCl, pH 6.5) was used for MMC and psoralen treatments under the following conditions: MMC at (1, 2.5, and 5) µg/mL MMC at 30 °C for 2 h; psoralen at (0.04 and 5 µg/mL) at 37 °C for 1 h, followed by 30 min UVA exposure; and psoralen at 0.04 µg/mL at 37 °C for 1 h, followed by 1 h of UVA exposure. An alternative Tris-Sucrose buffer (10 mmol/L Tris, 0.5 mol/L sucrose, pH 6.5) was also tested with psoralen under the same conditions as above. *Mcap*ΔRE recipient cells were grown in SSG medium at 30 °C to pH 6.7 ± 0.1. For each reaction, 3 mL of cells were collected, and MMC or psoralen were added to achieve the identified final concentrations. Treated cultures and their corresponding controls were incubated under the conditions described above. WGT was performed as described above, using centrifugation at 4575 × *g* and digested mycoplasma plugs. Following the final resuspension, reactions were plated directly on SP4 agar plates containing tetracycline without a recovery period.

### WGT on Selection-Free Plates

To assess whether antibiotic selection is necessary for selection-free WGT, WGT was performed on SP4 agar plates with and without 4 µg/mL tetracycline. Digested JCVI-syn1.0 bacterial and yeast plugs were used as donor DNA. *Mcap*ΔRE Recipient cells were grown in SSG medium at 30 °C to pH 6.7 ± 0.1. For each reaction, 3 mL of recipient cells were used for 20 µL mycoplasma plugs, and (3 or 9) mL of recipient cells were used for 100 µL yeast plugs.

MMC was added at a final concentration of 2.5 µg/mL or an equivalent volume of DMSO for controls, and then incubated at 30 °C for 2 h. Transplantation was performed as described above. Eight replicates were included for each condition mentioned above. After final resuspension, four reactions were plated on SP4 agar plates containing tetracycline and four reactions were plated on SP4 agar plates without tetracycline; there was no recovery period. For comparison, untreated recipient cells were plated on SP4 agar plates containing X-gal at a 10^6^ dilution after final resuspension.

### Multiplex PCR confirmation

After transplantation on selection-free plates, white and blue colonies were individually picked. Seven colonies (blue or white) were inoculated into 1 mL of SP4+KO medium with and without 4 µg/mL tetracycline in separate 1.8 mL tubes, for a total of 28 tubes, and incubated at 37 °C (8 to 16) h. 50 µL of cells from each tube were mixed with an equal volume of water and boiled at 95 °C for 5 min to release genomic DNA. 2 µL of the resulting lysate from each tube was used as the DNA template. Each template was resuspended in TE buffer to a final concentration of 2.0 µmol/L. A master mix was prepared containing nine amplicon pairs (listed in the Table 1, ordered from IDT), along with yeast rRNA (Amplicon size of 464 bp) and His3 (Amplicon size of 230 bp) primer pairs, for a total of 11 amplicons (22 primers). Multiplex PCR was performed using Multiplex PCR kit (Qiagen, 206143) with a 15 µL reaction containing 7.5 µL of 2 × Qiagen Multiplex PCR Master mix, 0.75 µL of 20 × primer mix (2.0 µM each), 2 µL of DNA template, and 4.75 µL distilled deionized water was used. This reaction was diluted to 40 µL with 25 µL TE buffer. Cycling conditions were initial heating at 95 °C for 15 min; 34 cycles of 94 °C for 30 s, 57 °C for 2.5 min, and 72 °C for 1.5 min; followed by 68 °C for 15 min. 20 µL of each reaction were analysed on a 1% E-gel (Invitrogen) by applying 72 V for 30 min, and bands were visualized using a Typhoon 9410 Imager.

## Supporting information

Supplementary Material: Selection-free whole genome transplantation revives dead microbes

## ACKNOWLEDGEMENTS

We acknowledge support from the U.S. National Science Foundation (MCB 1840320, MCB 1818344, MCB 1840301, and MCB 2221237), the U.S. National Institute of Allergy and Infectious Diseases (R21 AI098057-02), and the U.S. Department of Agriculture (NIFA 2015-67016-23169), and the Ruggles Family Foundation. We are also grateful to our collaborators and proud to contribute to the growing field of synthetic biology.

The authors declare no financial or commercial conflicts of interest. To the best of our knowledge, no financial relationships exist that have influenced the research described in this work.

Figures 1a was generated using BioRender (http://BioRender.com), which was employed to create schematic representations of the experimental workflow and key procedural steps.

## Disclaimer

Certain commercial entities, equipment, or materials may be identified in this document in order to describe an experimental procedure or concept adequately. Such identification is not intended to imply recommendation or endorsement by the National Institute of Standards and Technology, nor is it intended to imply that the entities, materials, or equipment are necessarily the best available for the purpose.

## Author Contributions

ZPS, NAG, VP, OC, FW, TG and JIG performed the experiments. EAS, ER, JCV and JIG provided critical feedback that guided the project direction. ZPS, NAG and FW analysed the experimental data, ZPS and NAG prepared the figures. ZPS, with input from VP and FW, drafted the initial manuscript. ZPS and JIG framed the discussion, and finalized the manuscript by integrating the text, figures, and references. JCV and JIG supervised the overall project. All authors contributed to manuscript revision, provided feedback, and approved the final version.

