## Supplementary Material: Selection-free whole genome transplantation revives dead microbes for "Selection-free whole genome transplantation revives dead microbes"

This material includes:

Methods and Results

Figures S1-S4

References

**Methods and Results**

**Microbial Strains**

*Mycoplasma genitalium* strains G37 and 1019V were obtained from the ATCC and as a gift from Joel Baseman at the University of Texas Medical Center at San Antonio, respectively*.* To add kanamycin and puromycin antibiotic resistance markers to *M. genitalium* G37 we constructed *E. coli* plasmids (pET Novagen), containing $\approx$2000 base pairs from either the MG408 or MG210 genes for kanamycin or puromycin resistance, respectively. Both plasmids lacked an origin of replication for mycoplasma. In two successive steps, the plasmids were chemically transformed into *M. genitalium* G37 and plated on nutrient agar containing the appropriate selective antibiotic. The M. genitalium homologous recombination system recombined the plasmid inserts into genes MG210 and MG408^1^. We confirmed the intended gene insertion using PCR amplification (data not shown).

**Intraspecies WGT in *M. genitalium***

We performed intraspecies WGT using donor G37^2^ and recipient 1019V strains. We built a G37 mutant with a puromycin antibiotic resistance marker inserted in gene MG210 and a kanamycin antibiotic resistance marker inserted in gene MG408. Thus, the markers were separated by more than 250,000 base pairs. We plated WGT reactions on SP4 agar containing 50 µg/mL kanamycin (Sigma-Aldrich PHR1487). The kanamycin colonies were then transferred onto SP4 agar containing 3µg/mL puromycin (Sigma-Aldrich P9620), and we observed no growth. None of the colonies were resistant to both antibiotics (data not shown). We concluded the kanamycin resistant colonies were false positives resulting from regions of the G37 genome containing the kanamycin marker exchanging with the corresponding region in the 1019V genome via homologous recombination.

**Pretreatment Duration, Temperature, and Wash Buffer Effect on WGT**

To determine conditions for MMC treatment for selection-free WGT, parameters including treatment duration, temperature, and wash buffer were evaluated. Recipient cells were grown to pH 6.7 ± 0.1. A wash buffer containing 10 mmol/L Tris-HCl and 25 mmol/L NaCl (pH 6.5) was tested at 4 °C for 2, 4 and 18 h, at 30 °C for 2 and 4 h, with 2.5 µg/mL MMC. We tested an alternate wash buffer of Tris-Sucrose buffer (10 mmol/L Tris, 0.5 mol/L sucrose, pH 6.5) for 2 h and 4 h incubations with 2.5 µg/mL MMC at 30 °C. Following incubation, transplantation was performed as described above with 3 mL recipient cells for each experiment, centrifugation at 4575 × *g,* and using digested mycoplasma plugs. Cells were plated on SP4 agar plates with tetracycline.

The results showed $\approx$50 % fewer transplants from MMC-treated cells relative to controls without MCC ($\approx($35 to 60 transplant colonies) versus $\approx($70 to 150 colonies) across incubation times or temperatures (Figure S1). Amongst the conditions tested, MMC in Tris-NaCl wash buffer supported comparatively higher WGT efficiency.

**WGT of an MMC-survivor colony**

To determine whether the viable cells remaining after MMC treatment (i.e., MMC survivor cells) exhibited enhanced transplantation efficiency, *Mcap*ΔRE recipient cells were grown until pH 6.7 ± 0.1. An MMC working solution (100 µg/mL) was prepared by diluting the MMC stock (10 mg/mL in DMSO) in SSG medium; a control solution was prepared similarly by adding an equivalent amount of DMSO to SSG. For each treatment, 50 µL of the corresponding MMC or control solution was added to 1 mL of cells, ensuring equal DMSO concentration across all samples. Cells were treated with 0, 1, 2.5 and 5 µg/mL MMC for 2 hours at 30 °C. After treatment, 300 µL of MMC-treated cells were plated onto non-selective SP4 agar plates (without tetracycline). Untreated cells were plated using 10 µL of 10^-6^ and 10^-7^ dilutions on non-selective SP4 plates. Plates were incubated at 37 °C for 2-3 days until colonies appeared. Single colonies from untreated and 2.5 µg/mL MMC-treated plates were then individually picked and grown at 30 °C until each culture reached pH 6.7 ± 0.1. Both untreated and MMC-treated cultures were divided into two groups: one receiving the control solution and the other exposed to 2.5 µg/mL MMC, maintaining identical DMSO concentrations. This yielded four experimental conditions. Transplantation was subsequently performed as described above, with centrifugation at 4000 × *g*, using digested mycoplasma plugs, and reactions were plated on tetracycline-containing SP4 agar plates following a 2 h recovery period.

The fact that the MMC concentrations that allow WGT do not kill all the recipient cells suggests that WGT may be taking place in the small fraction of the surviving *Mcap*ΔRE recipient cell. Figure 1e data indicated this is extremely unlikely. First, 2.5 µg/mL MMC treatment reduced the recipient cell concentration by a factor of 10^6^, but the number of transplants dropped only by a factor of 10. Thus, WGT without MMC yields one transplant per $\approx$1.5 × 10^8^ living recipient cells. By contrast, if WGT is only happening in the few remaining living cells, we produced one transplant per $\approx$285 viable cells. That would require a 500,000-fold increase in WGT efficiency and would suggest WGT is taking place in dead cells and/or the MMC resistant cells were “super transplanters.” We isolated the *Mcap*ΔRE recipient cell that survived MMC and tested them to see if they were more competent for WGT. They were not (data not shown). Additionally, Figure 2 we showed that while transformation of recipient cells with a plasmid capable of replicating in *M. capricolum* yielded hundreds of colonies, the same transformation of MMC treated cells yielded zero transformants. Further evidence the recipient cells were killed. Collectively these results demonstrate that at least some of the transplants are synthetic living cells made by the installation of synthetic DNA into dead cells.

**Figures**

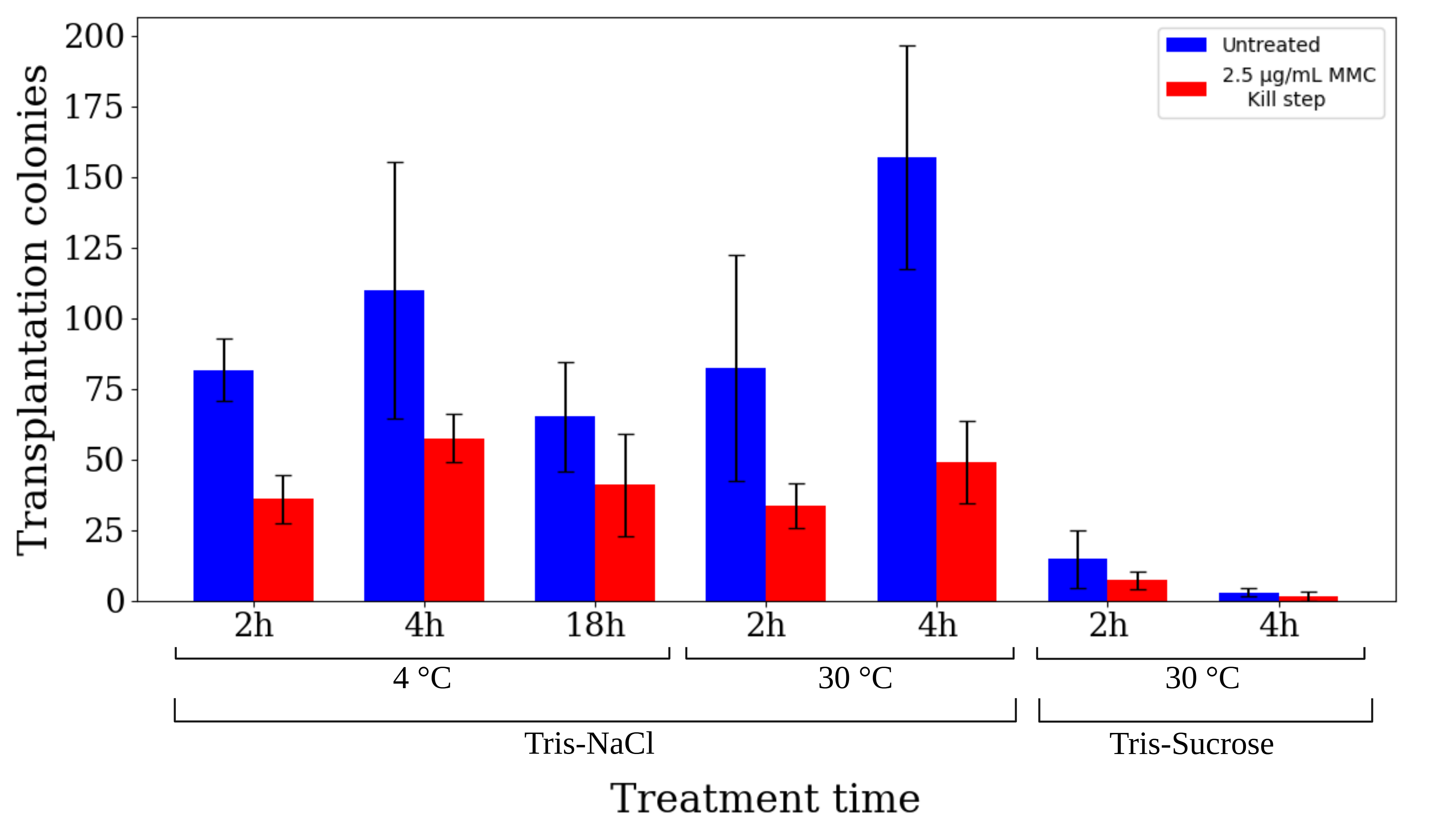

**Figure S1. Optimization of MMC pretreatment parameters for selection-free WGT. Treatment duration, incubation temperature, and wash buffer composition were evaluated for their effects on WGT.** The bar graph shows arithmetic mean values, and black error bars represent standard deviation from three biological replicates.

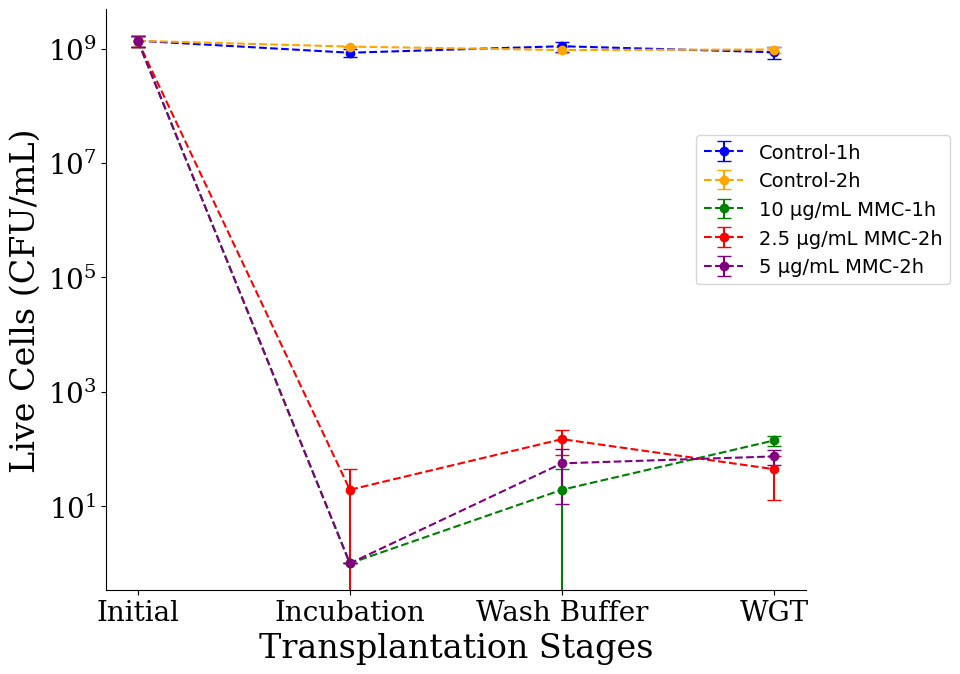

**Figure S2. Living recipient cell counts at different stages of WGT and selection-free WGT. MMC-treated recipient cells (Figure 2c and 2d) decline from 10^9^ CFU/mL to low hundreds, whereas untreated controls retain high viability.** Each data point represents arithmetic mean values of CFU plates, and error bars represent standard deviation from three biological replicates. Lines are drawn to guide the eye.

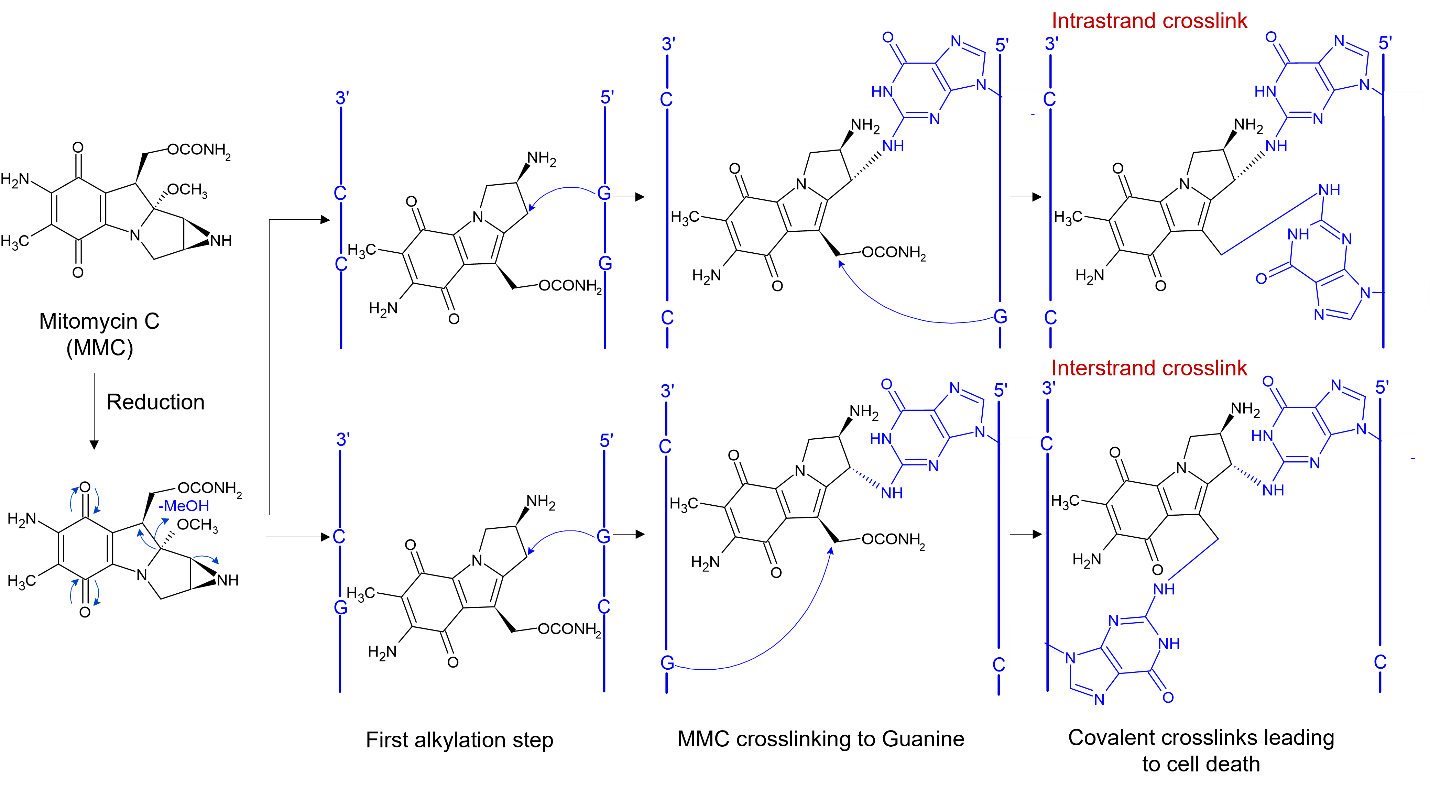

**Figure S3. Mechanism of MMC treated genome disruption through guanine nucleoside DNA crosslinking^3,4^.**

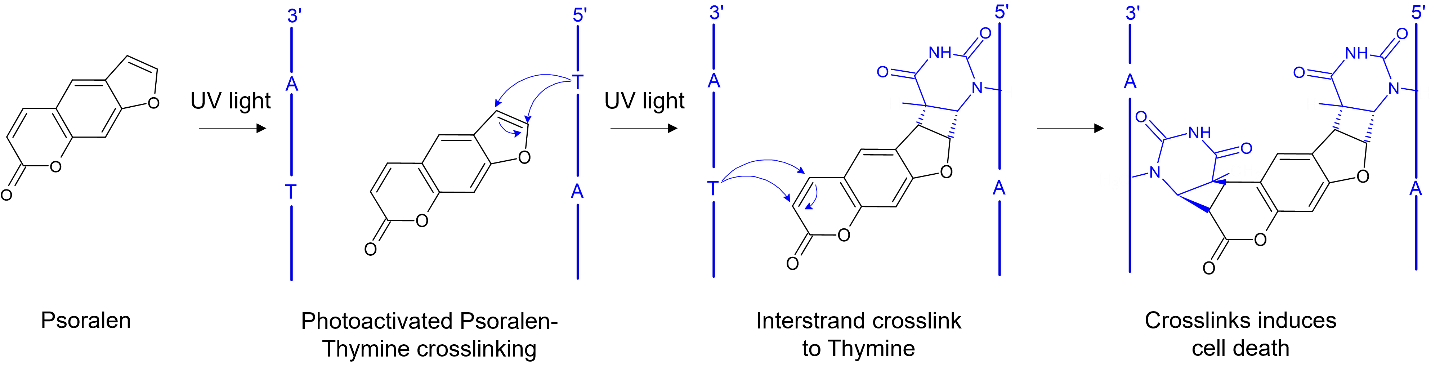

**Figure S4. Mechanism of psoralen treated DNA crosslinking triggered by light-induced (UVA) activation^5^.**
